# Unraveling protein interactions between the temperate virus Bam35 and its *Bacillus* host using an integrative Yeast two hybrid-High throughput sequencing approach

**DOI:** 10.1101/2021.05.11.443545

**Authors:** Ana Lechuga, Cédric Lood, Mónica Berjón-Otero, Alicia del Prado, Jeroen Wagemans, Vera van Noort, Rob Lavigne, Margarita Salas, Modesto Redrejo-Rodríguez

## Abstract

*Bacillus* virus Bam35 is the model *Betatectivirus* and member of the *Tectiviridae* family, which is composed of tailless, icosahedral, and membrane-containing bacteriophages. The interest in these viruses has greatly increased in recent years as they are thought to be an evolutionary link between diverse groups of prokaryotic and eukaryotic viruses. Additionally, betatectiviruses infect bacteria of the *Bacillus cereus* group, known for their applications in industry and notorious since it contains many pathogens. Here, we present the first protein-protein interactions network for a tectivirus-host system by studying the Bam35-*Bacillus thuringiensis* model using a novel approach that integrates the traditional yeast two-hybrid system and Illumina high-throughput sequencing. We generated and thoroughly analyzed a genomic library of Bam35’s host *B. thuringiensis* HER1410 and screened interactions with all the viral proteins using different combinations of bait-prey couples. In total, this screen resulted in the detection of over 4,000 potential interactions, of which 183 high-confidence interactions were defined as part of the core virus-host interactome. Overall, host metabolism proteins and peptidases are particularly enriched within the detected interactions, distinguishing this host-phage system from the other reported host-phage protein-protein interaction networks (PPIs). Our approach also suggests biological roles for several Bam35 proteins of unknown function, resulting in a better understanding of the Bam35-*B. thuringiensis* interaction at the molecular level.

**Author summary:** Members of the family *Tectiviridae*, composed of non-tailed icosahedral, membrane-containing bacteriophages, have been increasingly scrutinized in recent years for their possible role in the origin of dsDNA viruses. In particular, the genus *Betatectivirus* receives increased attention as these phages can infect clinical strains as well as industrially relevant members of the *B. cereus* group. However, little is known about the interactions between these temperate viruses and their hosts. Here, we present the first high-throughput study of tectivirus-host protein-protein interactions focusing on Bam35, model virus of betatectiviruses, and its host *B. thuringiensis*, an important entomopathogenic bacterium. We adapted the well-known technique yeast-two-hybrid and integrated high-throughput sequencing and bioinformatics for the downstream analysis of the results which enables large-scale analysis of protein-protein interactions. In total, 182 detected interactions show an enrichment in host metabolic proteins and peptidases, in contrast with the current knowledge on host-phage PPIs. Specific host-viral protein-protein interactions were also detected enabling us to propose functions for uncharacterized proteins.

## Introduction

The *Tectiviridae* family is defined as a family of tailless, icosahedral viruses with a lipidic inner membrane and a double-stranded linear DNA genome of approximately 15 kb, capped by the so-called terminal protein [1]. These phages are currently divided into three genera: (i) the alphatectiviruses, which are lytic phages infecting Gram-negative bacteria; (ii) the betactectiviruses, non-integrative temperate phages infecting Gram-positive bacteria [2]; and (iii) the gammatectiviruses, containing temperate phages that infect Gram-negative bacteria [3]. Together with the other recently discovered *Tectiviridae*, this family is characterized by well-conserved features, with a broad genetic variety and host-range, and is of ecological importance [4,5]. Additionally, they were proposed to be related to the origin of some mobile elements and groups of eukaryotic DNA viruses [6,7]. Among the *Tectiviridae* members, the interest on betatectiviruses has increased in the last decade due to their ability to infect different members of the *Bacillus cereus* group [8]. Indeed, some specifically infect pathogenic bacteria such as phages Wip1 and AP50 that can infect *B. anthracis*, the etiological agent of anthrax [9,10], or Sole and Simila that infect the food pathogen *B. cereus* [11].

The model virus for the molecular and structural studies on betatectiviruses is *Bacillus* virus Bam35 [12–15]. This phage infects *B. thuringiensis*, a type species of the *B. cereus* group, known for its entomocidal capacity and broadly used as biopesticide for pest control [16,17]. The betatectivirus genome organization is modular and traditionally segmented into three functional categories: (i) gene regulation & genome replication, (ii) virion structure & DNA packaging, and (iii) host recognition & cell lysis [18]. Despite the conserved genome organization between Bam35 and the widely characterized model virus for Alphatectiviruses phage PRD1, there is very low sequence identity between them. Moreover, the limited similarity to proteins published in databases limits functional annotation of Bam35 and other tectiviruses. This problem has been addressed by comparative studies, single protein purification and analysis, and protein-protein interaction (PPI) studies, resulting in the current functional annotation of 23 out of the 32 open reading frames of Bam35 [12,14,19,20].

The icosahedral capsid of Bam35 is mainly composed by the major capsid proteins that form the facets, and the penton proteins that are located in the eleven vertices and incorporate the flexible spikes. The packaging and injection of DNA takes place through the 12^th^ vertex, also known as the special vertex [13]. Both capsid proteins and dsDNA interact with the inner membrane lipids [13,21]. Although not all the membrane proteins have been identified for Bam35, P25 is probably a membrane structural component and P26 is a conserved transglycosylase that seems to be a cornerstone transmembrane protein that interacts with both lytic and capsid proteins [20,22].

Bam35 has been proposed to infect host cells following a three-step mechanism. First, the flexible spikes recognize and bind the cell surface receptor. So far, only one of the components of this receptor has been identified, the N-acetyl-muramic acid, which is essential for phage adsorption [23]. Second, the peptidoglycan hydrolyzing proteins facilitate overcoming the cell wall barrier to access the plasma membrane. Finally, as in PRD1, Bam35 forms a tail-like structure which consists of a proteo-lipidic tube that protrudes from the inner lipid membrane and delivers the linear dsDNA into the cell [13,23,24]. The Bam35 genome is replicated by a protein-primed mechanism which uses a so-called terminal protein (TP) to prime the genome replication and thus remains linked to the 5’ DNA ends [14,25,26]. Upon infection of the cell, this temperate phage can establish a lysogenic state as a linear episome. The lysis-lysogeny switch has been studied on GIL01 virus, which is almost identical to Bam35 [27]. During lysogeny, the host transcription repressor LexA remains bound to the viral protein P7 and restricts transcription of the late genes [19,28,29]. Although only low-expression levels have been suggested during the GIL01/Bam35 temperate phase, the phage nonetheless impacts bacterial growth, sporulation, motility, and biofilm formation [30]. Host cell activation of the SOS response allows the phage to enter the lytic cycle, which is mediated by the elimination of the transcription repression and the activation of the late genes transcription by the viral protein P6 [31]. In a final step during the lytic cycle, Bam35 virions are released by lysis, likely through an endolysin-holin system [32]. Although two endolysins have been described (P26 and P30), no holin has been identified to date [22].

The protein intraviral interactome of Bam35, as well as for other viruses, has recently contributed to unravel new functions and the localization of phage ORFan proteins [20,20,33]. However, understanding host-virus PPIs is also essential to study protein functions, life cycle, evolution, and is particularly helpful to identify molecular targets to combat pathogens [34]. The last decade has seen the emergence of “omics” approaches as investigative tools for the study of biological pathways involved in pathogen replication, host response, and, eventually, infection progression. Among the methods used in high-throughput interactomics, the yeast two-hybrid system (Y2H) remains one of the most widely used techniques to study PPIs [35]. Some of the first Y2H studies involving viruses addressed the interaction between bacteria and their phages. These include *E. coli* phages T7 and λ, *Pseudomonas aeruginosa* phages, *Streptococcus pneumoniae* phages Cp-1 and Dp-1, and mycobacteriophage Giles [36–41]. Other techniques for detecting PPIs, such as affinity purification coupled to mass spectrometry, have also significantly contributed to the study of phage-bacteria interactions [42,43]. These works suggest different and specialized PPI networks, reflecting their genetic diversity, distinct biology, and diverging co-evolutions with their specific hosts [44]. Despite phage-host specialization, proteins of phages infecting the same host are suggested to employ similar strategies. All of them appear to share a tendency to interact with central “hub” proteins, highlighting their potential disruptive effect to the host metabolism. Another commonality is found in the targeting of proteins involved in transcription, replication, recombination, and repair functions [40].

The recent combination of Y2H with high-throughput sequencing technologies (HTS) overcomes the labor-intensive clone-by-clone analysis and has been shown to speed up the study of PPIs, while increasing the efficiency and sensitivity of the method. Different approaches have been followed such as recombination-based methods [45–47] or methods based on the screen of a genomic library against one single protein [48,49]. Although these techniques represent a marked improvement of the method, the data analysis and interpretation remain a challenge. Indeed, the large amounts of data generated with these approaches require the development of specific bioinformatics pipelines, and a fine-tuning of thresholds to disclose reliable interactions.

To date, few large-scale analyses of phage-bacteria PPIs have been conducted. These works focus on model viruses from the *Caudovirales* order [37–41] and, to our knowledge, no detailed studies on phage-host PPIs have been performed on other groups of phages, including the *Tectiviridae* family. Besides, these existing studies rely on individual clones analysis of the Y2H screen results. In our work, we used a novel high-throughput approach, aiming to obtain an proteome-wide virus-host protein interactome between the betatectivirus Bam35 and its host, *B. thuringiensis*. By performing a total of 156 Y2H assays, we established a highly selective interactome Bam35-*B. thuringiensis* in which we could detect patterns within the phage-host interactions and identify specific interactions to better understand viral protein functions and phage biology.

## Results

### Integrating the yeast two-hybrid system with high-throughput sequencing

To obtain an extensive protein-protein interactome of Bam35 and *B. thuringiensis* through the development of a novel and customizable approach, we established an experimental setup that combines traditional Y2H with HTS. We used the previously generated Bam35 ORF collection [20], containing bait constructs for all 32 ORFs of Bam35, cloned in both orientations (C- and N-terminal fusions to the Gal4p DNA binding domain, DBD). This collection also includes truncated versions of proteins (labeled with a “t”) with a predicted transmembrane domain, from which this domain was removed. The Bam35 ORF constructs were used to screen two newly generated genomic libraries of *B. thuringiensis*.

The genomic libraries of the *Bacillus thuringiensis* HER1410 genome (6,147,475 bp) suitable for Y2H were obtained using a four-step procedure (Supplementary Figure 1). These libraries were generated by partial digestion of the genomic DNA followed by insertion of the obtained fragments into the Y2H prey expression vectors pGADCg and pGADT7g (pPC and pPN) respectively generating C-terminal and N-terminal fusions of DNA-binding Gal4p activation domain (AD) to the Gateway cassette.

To assess the quality of the genomic Y2H library in detail, an Illumina deep sequencing analysis was performed, using a custom bioinformatics pipeline (Figure 1). Selected fragments of the libraries containing ATG-starting reads (97% for pPC and 96% for pPN) were converted into HER1410 fragments that, after clustering, yielded at least 40,000 different HER1410 fragments for each library. Mapping of these unique fragments against the host genome resulted in a coverage of 40% of the total nucleotides of HER1410, almost identical for both libraries (40.8% for pPC and 40.07% for pPN). Also, both libraries are equally distributed throughout the genome with similar coverage (Supplementary Figure 2). Two additional filters were applied to the unique fragments based on their translation and position with respect to the *GAL4-AD* gene (the GAL4p activation domain-encoding part). Next, the clusters of unique fragments were reconstructed to obtain the frequency of each fragment. The final Y2H-validated fragments represent a total of 417 genes of HER1410 in the pPC and 781 genes in the pPN library (Supplementary Table 1). In this case, the difference in the number of represented genes by each library, lower for pPC, is mainly due to the differential final filtering step, which is more stringent for this combination.

**Figure 1.**
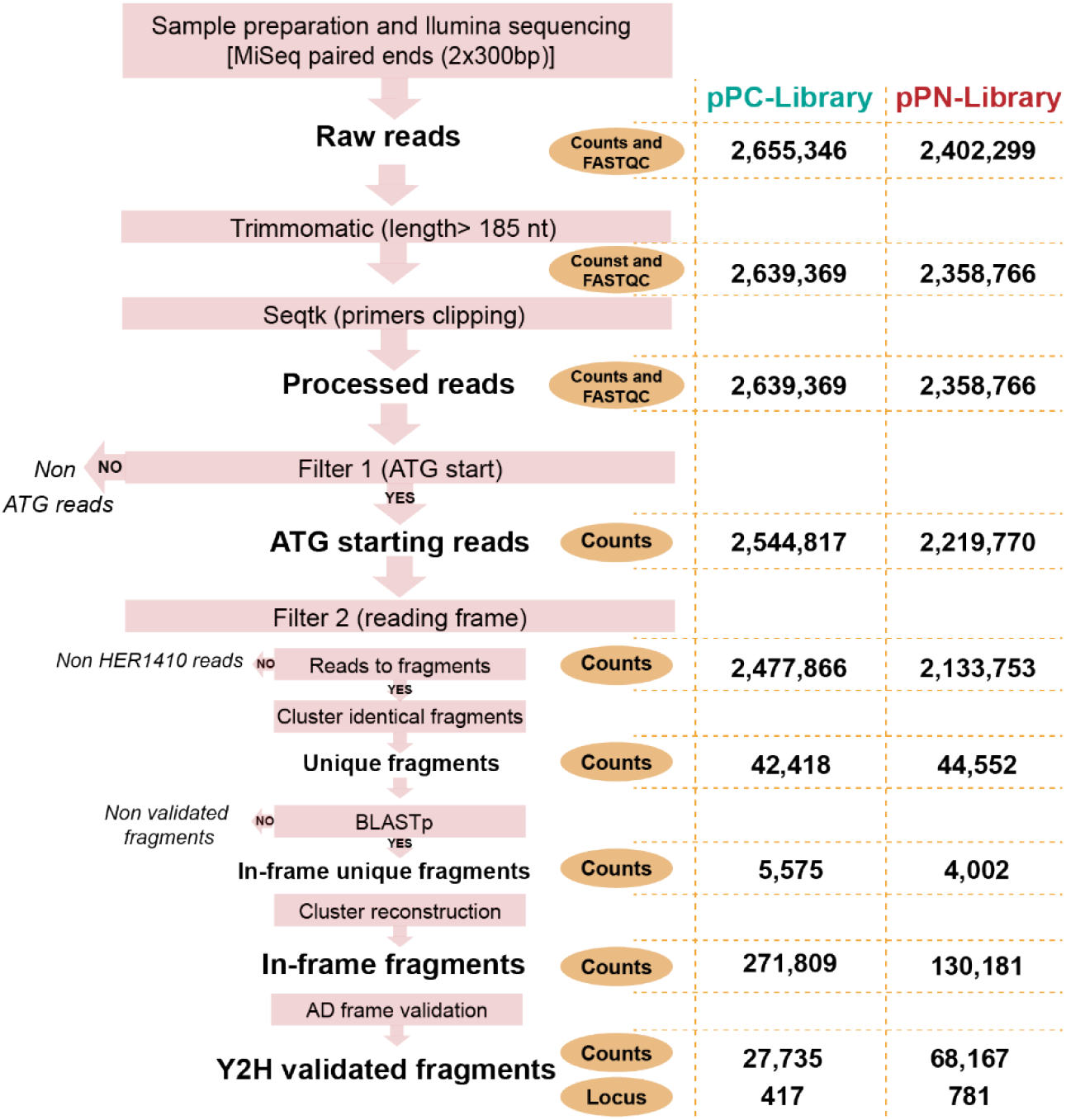
*B. thuringiensis* HER1410 genomic library high-throughput analysis. Bioinformatics pipeline for Y2H-HTS data analysis. Schematic representation of the different steps followed for sequencing data processing of Y2H libraries and, later on, Y2H positives (see next section). After each step, the number of reads or fragments was calculated (counts). These counts, for each library, are indicated in the two columns on the right (pPC: teal, pPN: red). In the first steps, FastQC was used to evaluate the quality of the reads, illustrated in Supplementary Figure 3.

The interaction analysis of all viral ORFs against the genomic library in both orientations would result in a total of 158 different possible combinations (Figure 2), including mating experiments of our libraries with the empty bait vectors, performed to minimize the number of false positives. However, given that the ORF16 of Bam35 was not available in the pPC vector [20], this number was reduced to 156 individual Y2H assays. For each assay, a bait yeast harboring the Bam35-ORF(X) bait expression vector was mated with one of the prey libraries. The overall calculated mating efficiency was 61.23%.

The positive interactions were first detected using the *HIS3* reporter gene by plating the mated cells on selective media lacking histidine (Figures 2 and 3A). To increase the stringency of selection when using the *HIS3* reporter gene, the autoactivation of the bait proteins was avoided by adding the optimal concentration of the inhibitor 3-amino-1,2,4-triazole (3AT) for each bait construct. Besides, the reduction of cell background and, therefore, the efficient growth and selection of positive colonies was achieved by two steps of replica plating (Figure 3A). As can be observed in Figure 3A, the screen led to a wide variety of results in terms of color and number of colonies. The assays that included the pPN-Library and were plated on selective media with low 3AT concentration (0-0.1 mM) resulted in a pink lawn (Figure 3A, lane 4). This could correspond to a false positive background caused by the pPN-Library which prevents the recovery of positive colonies. For these cases, the increase of 3AT concentration up to 3 mM effectively avoided the pink lawn, allowing the growth of hidden interacting partners (lane 5). In addition, based on the *MEL1* reporter selection system, 5-Bromo-4-chloro-3-indolyl α-D-galactopyranoside (X-α-Gal) was added to identify and amplify reliable interactions by increasing the 3AT concentration in the plates containing mostly white colonies (weak interaction). However, the plates meeting this requirement already contained the highest concentration of 3AT. To identify the prey interaction partners in each assay, cells from the last replica plate (Y2H positive clones) were pool-harvested and the prey fragments were amplified by PCR for subsequent Illumina sequencing (Figure 3B). The PCR products of each assay showed distinct patterns of discrete bands or a smear which corresponds to different sizes within the library size range. Nine combinations showed few or no colonies and, consequently, no PCR product was generated (Supplementary Table 2).

**Figure 2.**
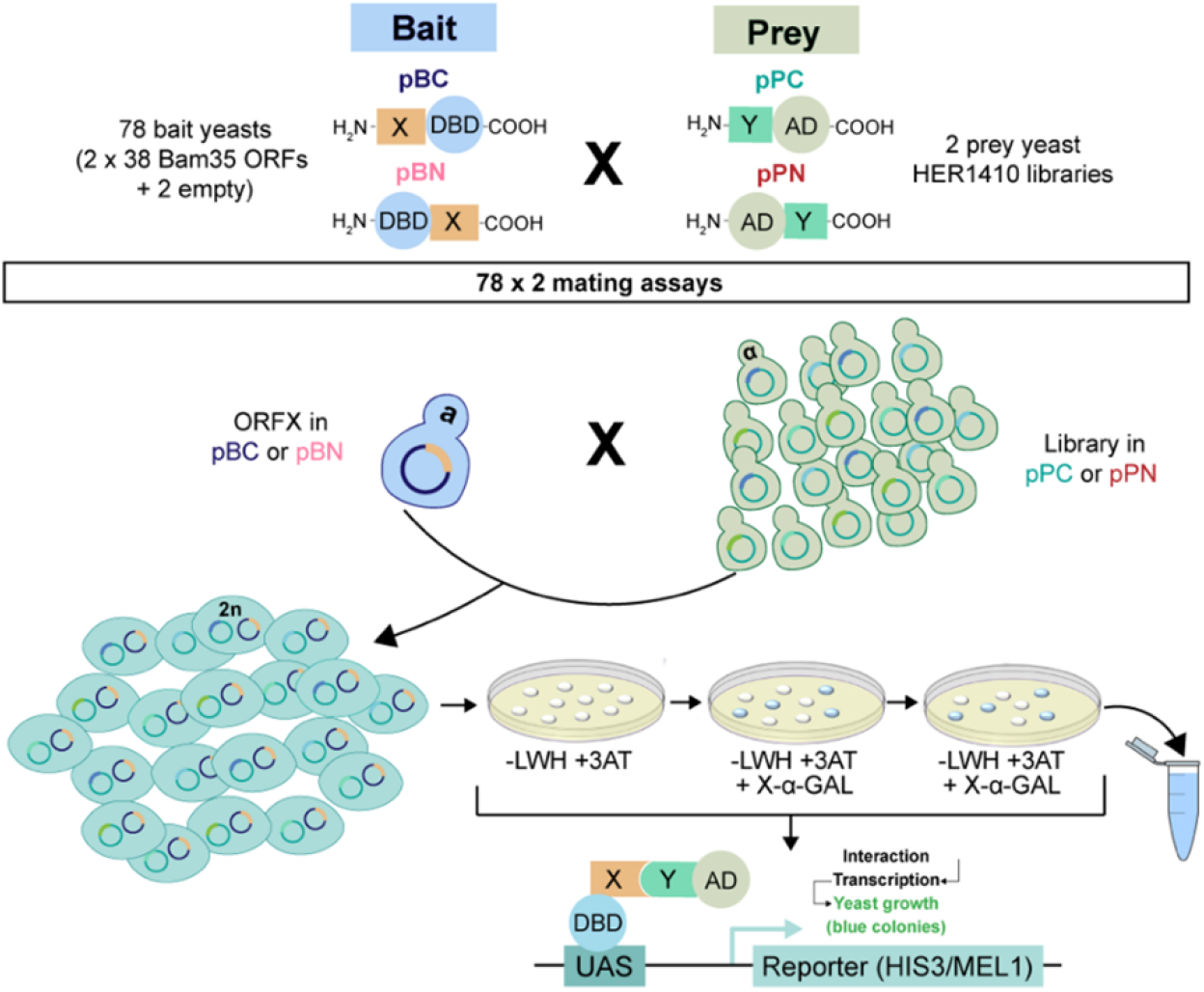
Bam35-*B. thuringiensis* Y2H interactome screen. Schematic representation of the Y2H assays. A total of 156 Y2H experiments were performed to test the interactions between each of the ORFs of Bam35 (X) cloned into bait vectors (C- and N-terminal fusion) and the generated HER1410 libraries (Y) cloned into prey vectors (C- and N-terminal fusion). For each assay, a Bam35 ORF-containing bait yeast was mated with one of the HER1410 library-containing prey yeast. Positive interactions were detected using *HIS3* and *MEL1* markers. Thus, mated cells were plated on selective solid media without leucine, tryptophan, and histidine (-LWH) supplemented with the corresponding 3AT concentration and then replicated twice in −LWH +3AT plates supplemented with X-α-Gal. The cells able to express the *HIS3* gene grew on −LWH plates and, further, those able to express *MEL1* turned blue in presence of X-α-Gal. All cells of the last replica were pooled and harvested for further analysis.

**Figure 3.**
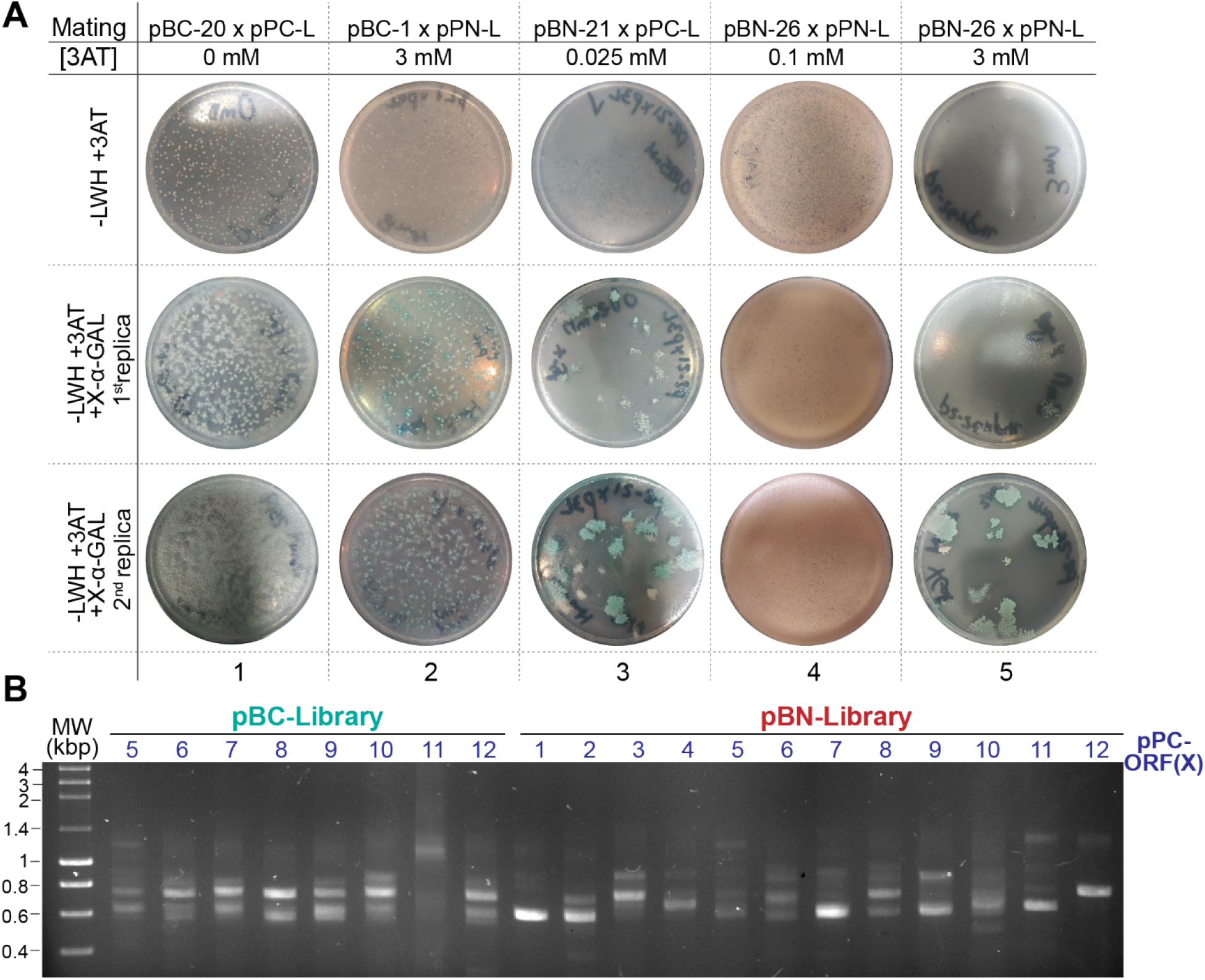
Representative output of the Bam35-*B. thuringiensis* Y2H screen. **(A)** Representative pictures of the three-step plating of four different bait-prey combinations. Plates from columns 4 and 5 correspond to the same combination with different 3AT concentration. Cells harboring interacting partners are able to grow in the absence of histidine, and therefore on −LWH media. Strong interactions were detected as blue colonies as a result of the metabolism of X-α-Gal in the first and second replicas. pBC-X and pBN-X stand for pBC and pBN harboring the corresponding viral ORF (X) and pPC-L and pPN-L stand for pPC and pPN HER1410 Libraries respectively. **(B)** Representative gel of the prey inserts PCR-amplification from different bait-prey matings. Cells from the last replica were harvested, their plasmid DNA was extracted, and the prey fragments were PCR amplified as explained in Materials and Methods to avoid amplification bias. For each sample/assay, several bands were detected whose size range is in agreement with the libraries size range.

### Data treatment reveals a good performance of the Y2H-HTS approach and generates a highly selective interactions dataset

The high-throughput analysis of each Y2H assay using Illumina sequencing (300 bp-paired-end run) resulted in 12,229 reads on average per sample (Supplementary Table 2). These reads were filtered using the bioinformatics pipeline illustrated in Figure 1. After data treatment, mapping of the Y2H-validated fragments allowed us to retrieve a total of 4,477 possible interactions (Table 1, Supplementary Table 3). A detailed analysis of these results, described in the Supplementary material, revealed that the Y2H screening strongly favors fragments in frame with the *GAL4-AD* gene and the actual ORFs within the genome, with only minor library-borne background that seems higher for the combinations including the pPN-library. The 4,477 interactions were classified into categories, according to their enrichment or presence in the Illumina reads dataset, as follows: A (100-10% of normalized counts), B (10-0.25%), and C (0.25-0%). These categories were validated by a small-scale sequencing analysis which showed a 100% recovery of prey fragments by HTS (with the exception of IS*4* transposases) and good correlation between their presence in the sample and their enrichment category (see Supplementary material for details).

**Table 1.** Frequency of interactions for each enrichment category. The table shows the total number of interactions and the proportion of sequencing reads (counts) that fall in each enrichment category.

| Enrichment |  | Number of interactions |  |  |  |  | Read counts (%) |  |  |  |  |
| --- | --- | --- | --- | --- | --- | --- | --- | --- | --- | --- | --- |
|  |  | pBC |  | pBN |  | Total | pBC |  | pBN |  | Total |
| Category | Range | pPC-Library | pPN-Library | pPC-Library | pPN-Library |  | pPC-Library | pPN-Library | pPC-Library | pPN-Library |  |
| A | 10-100 % | 93 | 43 | 67 | 27 | 230 | 77.89 | 72.61 | 73.75 | 81.09 | 75.93 |
| B | 0.25-10% | 262 | 165 | 391 | 95 | 913 | 20.76 | 24.34 | 24.52 | 15.76 | 22.03 |
| C | 0-0.25% | 738 | 794 | 990 | 812 | 3,334 | 1.35 | 3.05 | 1.73 | 3.15 | 2.04 |
| Total |  | 1,093 | 1,002 | 1,448 | 934 | 4,477 | 100 | 100 | 100 | 100 | 100 |

A similar number of total interactions, around 1,000, were found for each of the bait-prey combinations CC, CN, and NN (pBC-pPC, pBC-pPN and pBN-pPN, respectively), although their distribution into categories varies. Thus, the major contributor to the category A interactions is the CC combination while the NN combination generated the lowest number of A interactions. On the other hand, the NC (pBN-pPC) combination resulted in one third plus more interactions than the other combinations contributing more to B and C total interactions but similarly to the category A. Overall, compared to the pPC-Library interactions, those involving the pPN-Library showed a higher percentage of reads in the category C.

Remarkably, although a total of 3,334 interactions out of 4,477 were grouped in the category C, they correspond to only 2.04% of the total reads (Table 1). These hits cannot be properly discriminated from the data noise (see Supplementary material) and were excluded from the main results to increase the specificity, resulting in a dataset of 1,143 interactions (Table 2). Besides, Y2H methods may have a high rate of false positives [50]. In addition to the use of the appropriate 3AT concentration to prevent the autoactivation of the baits, two data filters were used to remove false positives. First, potential “sticky” preys, i.e, promiscuous prey fragments that interact with more than the average number of different bait interactors (six), were identified. A total of 36 “sticky” preys (Supplementary Table 4) were removed, reducing the dataset to a total of 228 filtered interactions (Table 2). Second, the prey proteins identified in diploid yeast harboring the empty plasmid (Supplementary Table 5) were also tagged as false positives and deleted from the dataset. Due to their tag as “sticky” preys, most of the prey fragments detected in the empty combinations had already been removed with the previous filter. Therefore, this second filtering step resulted in a reduction of only 17 hits.

**Table 2.** Total number of interactions detected by Y2H screening after each filtering step.

| Combination | Total | A-B category | No “sticky” preys | No empty | No duplicates |
| --- | --- | --- | --- | --- | --- |
| pBC-pPC_Library | 1,093 | 355 | 31 | 25 | 15 |
| pBC-pPN_Library | 1,002 | 208 | 24 | 15 | 14 |
| pBN-pPC_Library | 1,448 | 458 | 119 | 119 | 106 |
| pBN-pPN_Library | 934 | 122 | 54 | 52 | 47 |
| Total | 4,477 | 1,143 | 228 | 211 | 182 |

Lastly, after the consolidation of the duplicated results, a final high-quality dataset of 182 PPIs was obtained (Supplementary Table 6). More than half of these interactions (106) were found with the NC bait-prey pair. The second pair retrieving more filter-passing interactions was the NN pair which suggests a higher degree of detection of putative interactions for the N bait fusion. A total of 13 interactions were detected in two different bait-prey pairs, and, interestingly, seven of them involved the Bam35 membrane structural component P25 (Supplementary Table 6).

### Challenging Y2H single hits from the fragment genomic library by ORFs pairwise Y2H assays

To further validate the detected interactions from the Y2H screen, selected using the computational analysis of the Illumina dataset, 33 randomly chosen interactions were re-evaluated (Table 3). In this case, binary Y2H was used to individually test the interactions between the complete host proteins and their putative viral partners. Prey vectors containing the selected HER1410 ORFs were obtained and assayed with the correspondent bait vectors, resulting in seven confirmed interactions (Supplementary Figure 4). On the other hand, twelve of them could not be confirmed since, despite the yeast expressing both Bam35 and *B. thuringiensis* proteins was able to grow, the maximum 3AT tolerated was equal or smaller than that of the yeast expressing only the prey or bait protein (Table 3). Strikingly, only NC interactions could be confirmed by this method, coinciding with the bait-prey combination more commonly found within the HTS predicted interactions. Most of the confirmed interactions were detected at high 3AT concentration which indicates a strong interaction.

**Table 3.**
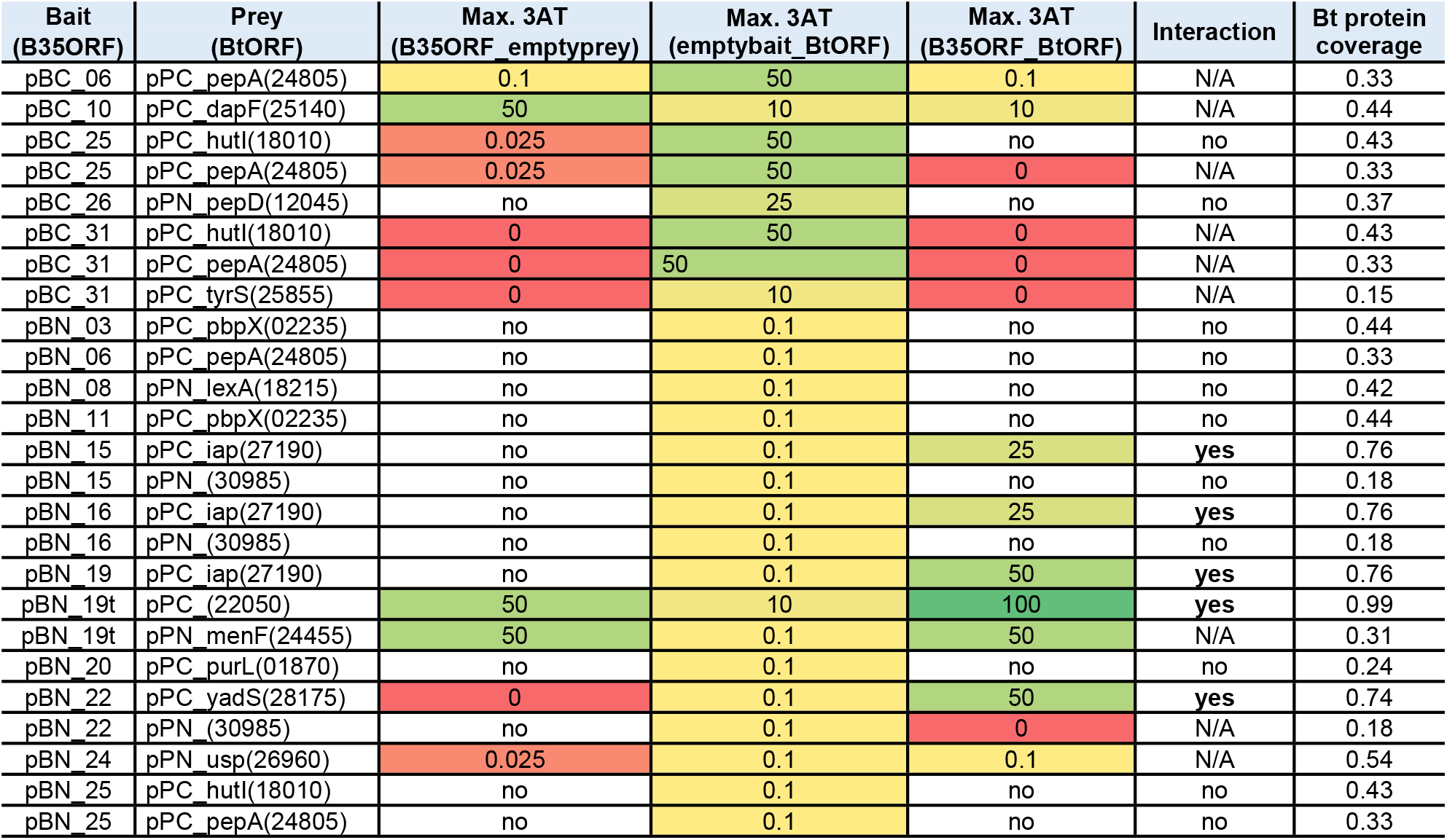
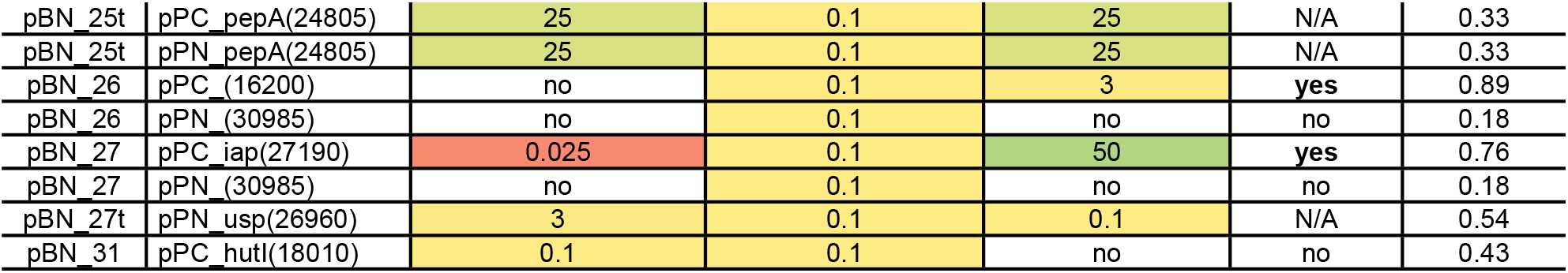
Binary Y2H screening of 33 selected putative interactions between Bam35 (B35) baits and *B. thuringiensis* (Bt) preys using full-length proteins. The maximum 3AT concentration at which yeast growth was detected is gradually colored from red (low concentration) to green (high concentration).

In this case, rather than the enrichment category, the percentage of the protein covered by the original library fragment resulted as a good indicator for the correlation fragment-full-length confirmation rate. Thus, all the interactions showing a host protein coverage above 74% were validated, while the rest (coverage under 54%) could not be validated for the full-length protein (Table 3). On the other hand, although only 21% of the retested interactions could be confirmed with this method, we cannot rule out that true interactions between viral proteins and host protein fragments or domains, which may be hidden within the full-length construct, occur. In conclusion, the confirmed positives rate provides a high confidence level for the interactions detected in our screening, at least for those involving high coverage fragments. These results suggest that the protein coverage of these fragments could also be useful as an additional confidence score to link protein fragment interactions with full-length PPIs.

### Bam35-*B. thuringiensis* Y2H predicted interactome: Clear the forest to predict PPIs

From the final dataset including the 182 seletected interactions, 54% include viral proteins functionally linked to the ‘Virion structure & DNA packaging’ functional group (Figure 4). Among them, both P26 and its non-transmembrane domain variant (P26t) retrieved the highest number of interactions (17 and 15 respectively), followed by P25 with 11 interactions (Table 4). In general, host metabolic proteins seem to interact to a higher extent with viral proteins than the rest of the clusters of orthologous groups (COGs), with more than half of the total PPIs including this type of proteins (Figure 4).

**Figure 4.**
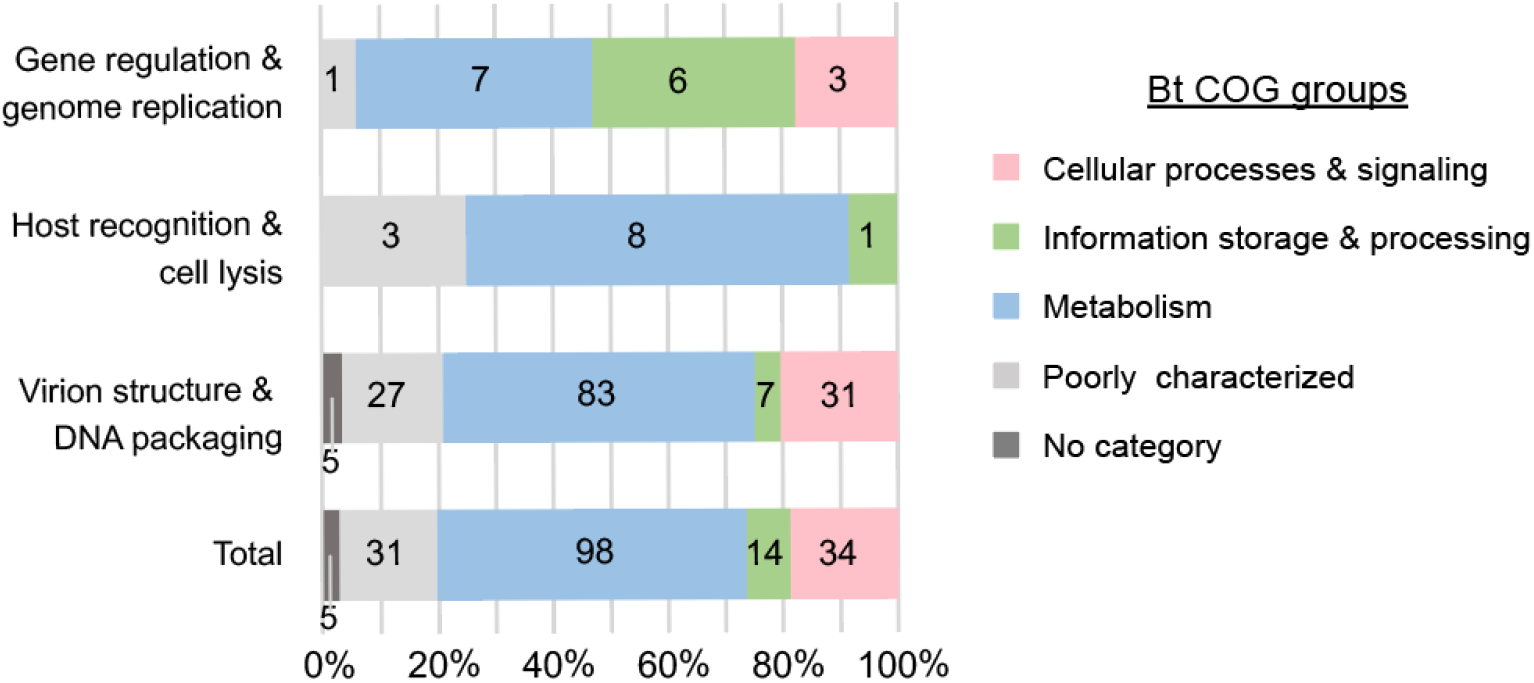
Interactions among functional groups of proteins. Stacked bar chart representing the proportion of interactions among the functional groups of Bam35 and the COG groups of *B. thuringiensis*. The number of total interactions is indicated for each combination inside the bars.

**Table 4.**
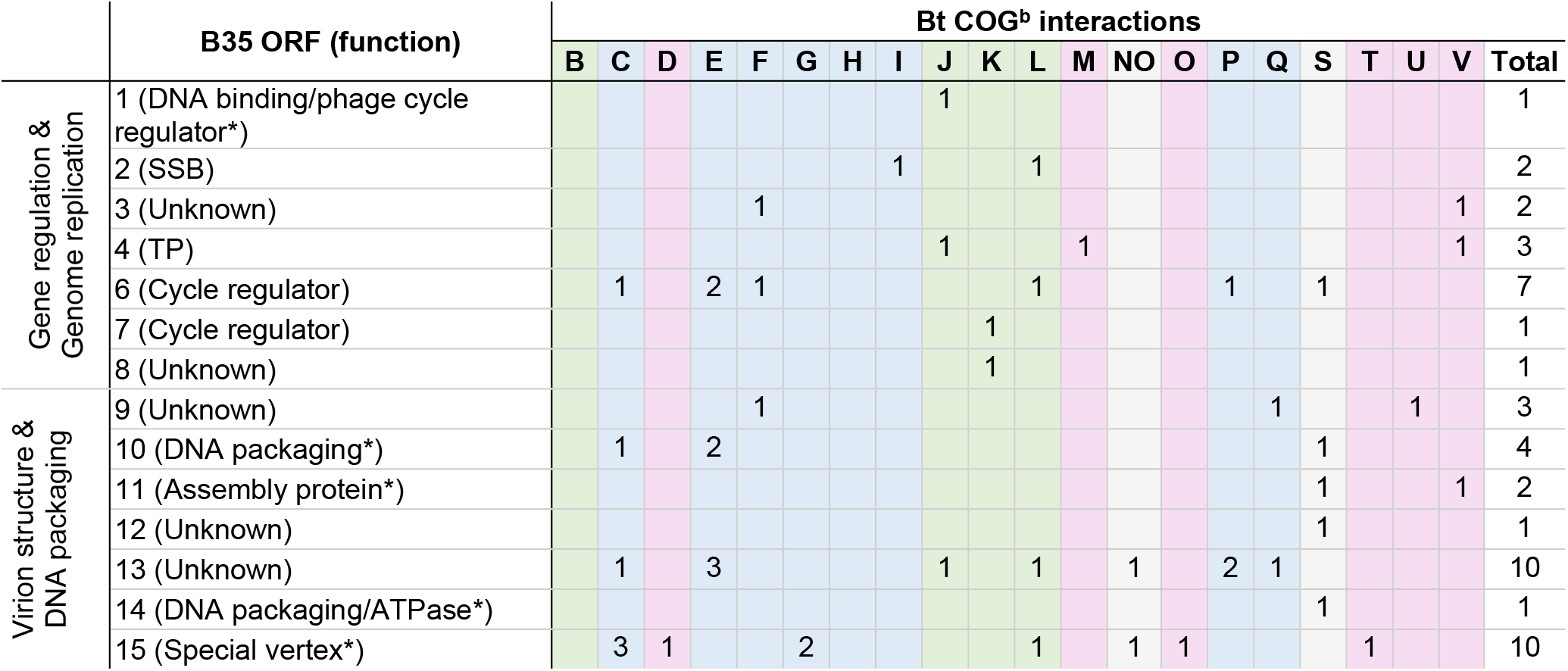
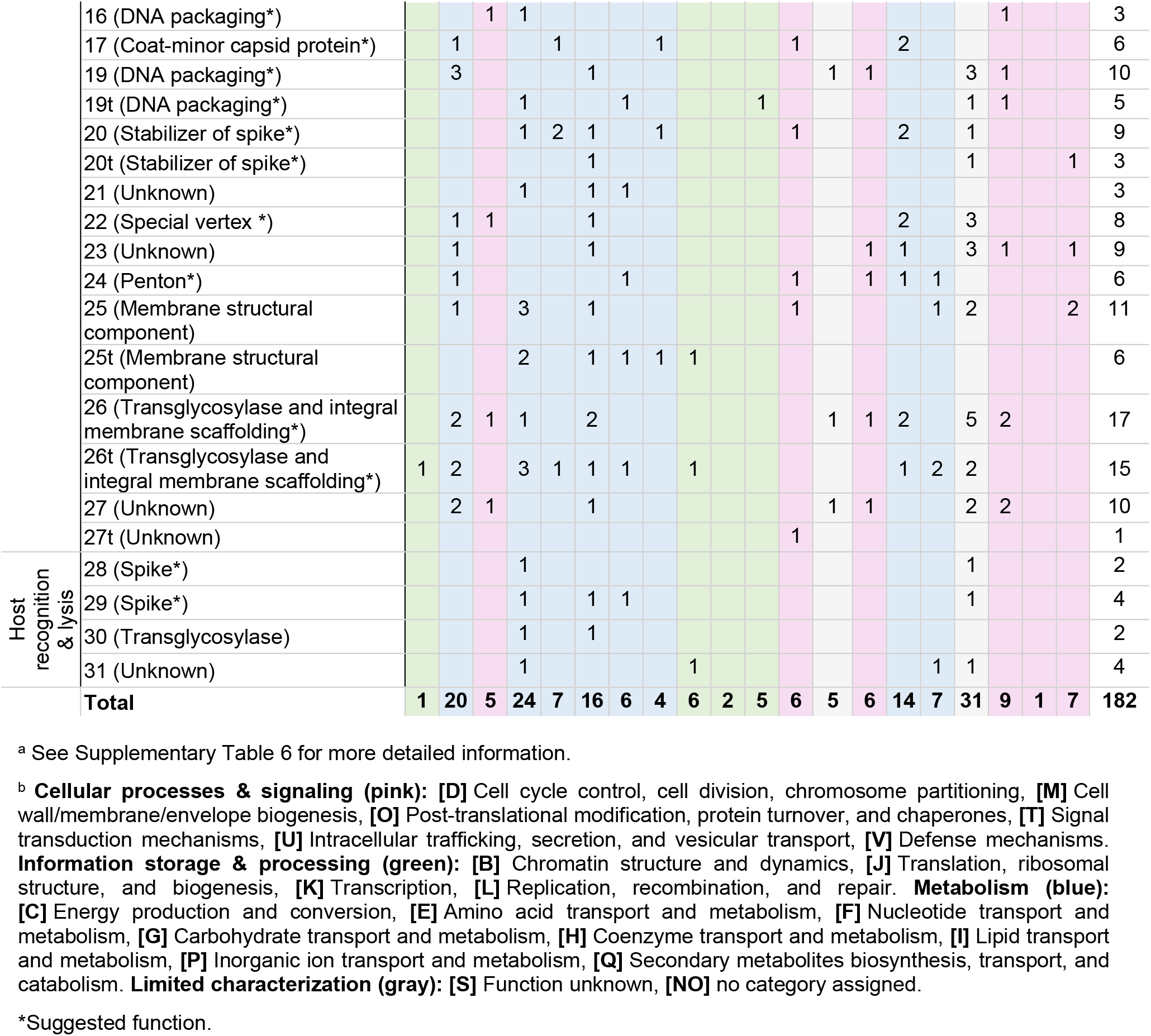
Putative PPIs between Bam35 (B35) proteins and *B. thuringiensis* (Bt) functional groups ^a^.

Interestingly, viral proteins involved in ‘Gene regulation & genome replication’ target ‘Information storage & processing’ host proteins in a higher proportion compared to other viral groups, indicating a possible link between these two functional groups. As shown in Supplementary Figure 5, those host partners do not connect different viral functional groups but remain restricted to replication and regulation viral nodes. The different viral functional groups of proteins seem to be connected mostly by metabolism host proteins. However, no connection was observed between ‘Host recognition & cell lysis’ and ‘Gene regulation & genome replication’ groups and only one node was linking the three groups of viral proteins. This interactor could be indeed a hub due to its role as aminopeptidase, a cytosolic protein presumably involved in the processing and regular turnover of intracellular proteins [51].

Analysis of the Bam35-*B*.*thuringiensis* interactome also allowed the identification of specific patterns and remarkable interactions (Figure 5 and Supplementary Table 6). Importantly, the only previously characterized Bam35-*B. thuringiensis* PPI, which consists of the interaction between the viral P7 and the host LexA protein [29], was detected in our final dataset. Interestingly, according to our results, this host protein could also interact with the viral protein P8, a protein of unknown function which, similar to P7, belongs to the ‘Gene regulation & genome replication’ functional group. Moreover, as explained above, the non-transmembrane variant of the protein P26 (P26t) appears to be a hub in the interactome since several host interactors were identified which also interact with other viral proteins. Also, the viral membrane protein P25 appears to be a structural hub that partners with several transport proteins. On the other hand, in agreement with the intraviral interactome results [20], the complete protein P26 interacts with host proteins that are also linked to several structural proteins, especially those involved in the special vertex formation (P15, P16, P19, P22), as well as P23 and P27 which are unknown function structural proteins. Therefore, these proteins seem to form a cluster connected by a high number of host proteins, many of them linked to membrane related transporters (Supplementary Table 6). Importantly, this “special vertex cluster” contains several high-confidence interactions including validated interactions, category A interactions and fragments that highly cover the complete host proteins.

**Figure 5.**
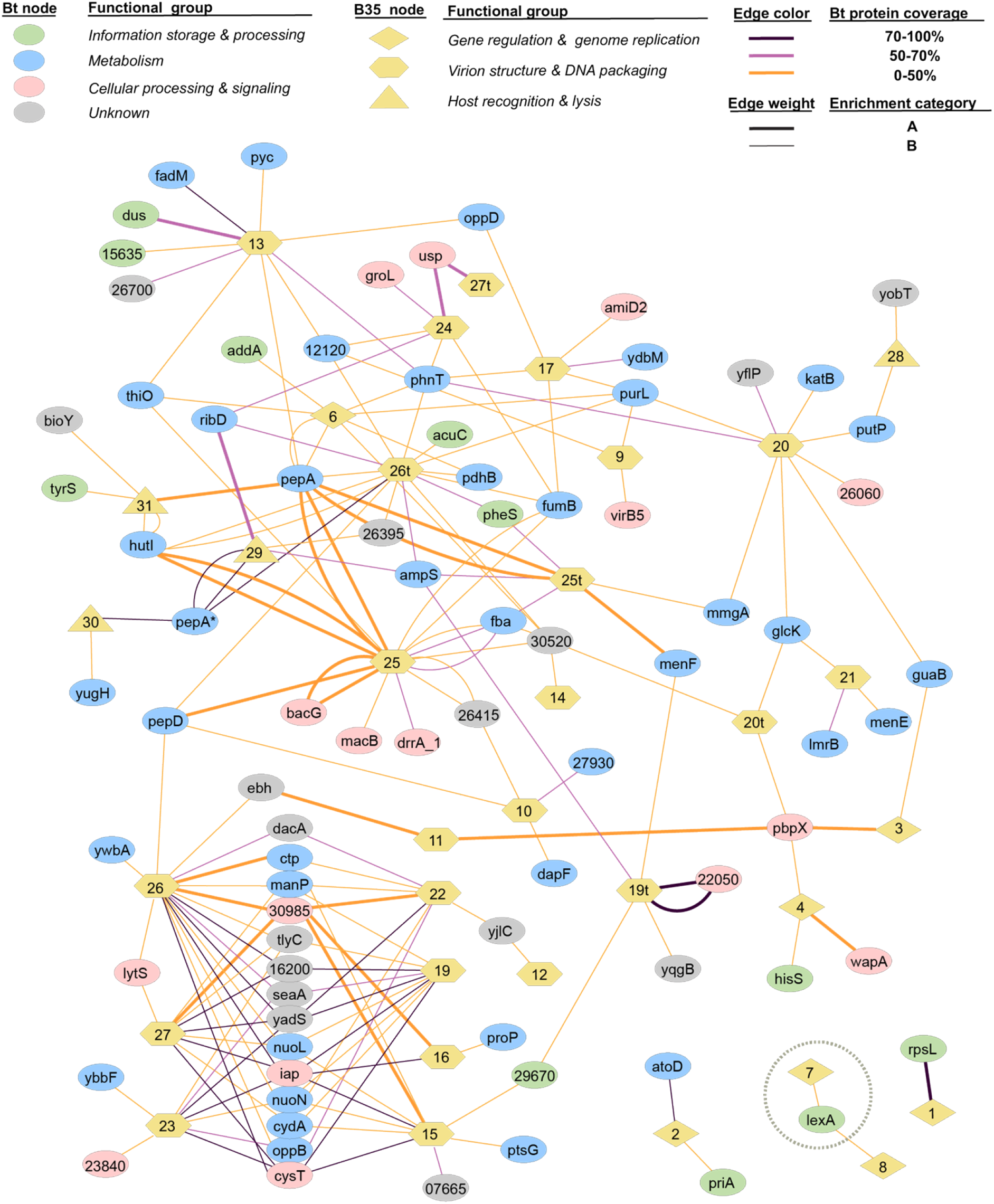
Bam35-*B. thuringiensis* protein interaction network based on the Y2H-Illumina screening. Network of the 182 detected interactions (195 bait-prey pair combinations) between the host proteins and the viral proteins (Supplementary Table 6). Host proteins (Bt nodes) are represented by ellipses colored according to their functional groups and the viral proteins are represented by yellow nodes shaped according to their functional groups. Each interaction (bait-prey combination) is represented as a line connecting two nodes. Line colors indicate the coverage of the host protein by the detected prey fragment and line weight indicates the enrichment category of the interaction (B for prey proteins detected as 0.25-10% of the sample counts and A for 10-100%). The interaction between the viral P7 and the host LexA detected in this work and previously published in Caveney *et al*. [29] is indicated with a gray circle. PepA* refers to the M42 family metallopeptidase encoded by the host gene HBA75_23575 and PepA refers to the cytosol aminopeptidase encoded by the host gene HBA75_24805.

Contrary to structural viral proteins that are very interconnected through host proteins, those involved in ‘Gene regulation & genome replication’ interact independently. This is the case, for example, of the B35SSB (P2) [52] which remains outside of the network, interacting only with an Acetate coA transferase and a primosomal protein. In general, the obtained network shows a highly interconnected system where some viral proteins could act as hubs in the bacteria-virus interactome [53]. Lastly, no interactions were detected between the host proteins and the viral P5 (DNA polymerase), and P18 (major capsid protein), which was somewhat expected by their function.

## Discussion

### A proficient Y2H-HTS method to detect multiple phage-host PPIs

To date, few attempts have been made to obtain bacteria-phage interactomes. Among them, the most complete includes an ORF collection of 3,974 (94%) of the known *E. coli* K-12 ORFs and 68 out of 73 ORFs of phage lambda whose interactions were tested by a pool-arrayed screening approach [38,54]. This method was also used in the case of *Streptococcus pneumoniae* and its phages, although with a less representative host ORFeome [40]. Although theoretically exhaustive, this approach requires automatization capacity and/or significant labor. It also hardly translates to less characterized models as it depends on the construction of an ORFeome. Others have turned to fragments-based approaches resembling this work for their PPIs analysis, like those of *Pseudomonas* phage ϕKMV and that of mycobacteriophage Giles [37,41]. However, both analyses of the libraries and the positives were based on individual clone identification, which negatively impacts efficiency, cost and labor. These limitations prompted us to use a combination of a random fragment library, and a novel pipeline which allowed us to benefit from high-throughput sequencing to study the interactome of Bam35 and its host, *B. thuringiensis*, and which could easily be portable to other systems. Coupling Y2H and HTS can be mainly performed using two strategies. An all-versus-all strategy using libraries recombination approaches or the use of barcode indexing which enables simultaneous sequencing of interacting preys from multiple separate assays in a single Illumina paired-end run [46,47,49,55]. In our case, the latter proved more appropriate since Bam35 has a limited number of viral genes, that were already available and tested for autoactivation in Y2H [20]. We also generated and thoroughly analyzed a random fragments genomic library of *B. thuringiensis* HER1410, a well characterized strain at the genomic level which is entomopathogenic and highly sensitive to *Bacillus* phage infections [18,56]. Based on this, our custom HER1410 library and the 32 encoded proteins from phage Bam35 have been screened using Y2H. This resulted in a large and high-throughput screening comprising 156 assays which potentially test more than 80,000 protein-protein pairs. A total of 4,477 interactions were initially detected, which were filtered and ranked to establish a threshold that differentiates low confidence interactions (or background) from reliable interactions. In our work, we showed that this approach is highly selective for in-frame fragments whose presence in the original library is at 1-2%, in line with previous work [57]. Indeed, some prey fragments from the Y2H positive results were not found in the libraries’ sequencing data. This suggests that these preys were scarce within the libraries and could not be detected despite the high sequencing depth of the library (over 200x). These results indicate that a higher depth could reveal the presence of additional fragments, increasing the calculated quality of the library. The establishment of categories based on the enrichment of the prey fragments per sample showed to keep one fourth of the interactions while retaining 98% of the Illumina reads (Table 1). Additionally, the validation of these categories by single-colony sequencing confirmed the strong correlation between the number of positive colonies and their enrichment in the Illumina data. This provides a quantitative value to the dataset, with reduced bias and high sensitivity, as previously proposed [48,49].

It is widely recognized that the main limitation of the Y2H is the high rate of false positives and false negatives. Caufield *et al*. [58] showed that the false-negative rate can be considerably reduced through the use of different vector pairs. Particularly, permutations of C- and N-terminal Y2H vectors increase the coverage of interactome studies, reduce the number of false negatives, and detect strong interactions [59]. However, for bacterial and phage-bacteria interactomes that include ORF or genomic libraries, those libraries are usually cloned in the N-fused variant for both preys and baits, possibly due to the restrictions imposed by the difficulty of creating “in-frame” C-terminal fusions [37,40,41,60,61]. In this work, the C-terminal fusion library gave rise to a high number of putative interactions when combined with the N-fusion viral proteins, despite harboring less in-frame fragments. This contrasts with some intraviral bacteriophage interactomes in which the NN combination appeared as the most frequent [20,58]. In addition, the pairwise screen using full-length host proteins only confirmed interactions that involve the C-fused library against the N-fused viral proteins. A total of 13 out of 182 putative interactions were found in more than one combination, showing a lower overlap than for intraviral interactomes [20].

The use of HTS to identify all the possible prey partners involved in the Y2H screening also reduces the false-negative rate to the minimum that Y2H can attain. We also used several methods aiming to maximally reduce false-positive interactions such as the competitive inhibitor 3AT which prevents bait self-activation, the addition of negative controls including the empty plasmids, and the identification and removal of “sticky” preys. Remarkably, a large percentage (97%) of the prey proteins found in the empty combinations were also tagged as “sticky”. Importantly, the high-throughput technique used in this work allows the identification of all prey partners, enabling the effective discrimination of specific interactors from “sticky” proteins and other false positives, as suggested elsewhere [49,62]. After the efficient removal of a high number of false positives (85%) by these filters, 182 candidate interactions could be identified.

The putative binary interactions detected in Y2H are often validated by other methods such as pull-down, co-immunoprecipitation, and surface plasmon resonance [37,63]. However, validation of high-throughput studies results is hardly feasible. As such, in line with our results, for large-scale methods computational biology and confidence scores are useful [49,64,65]. In addition, Y2H approaches generally validate their methods by retesting the interactions with pairwise Y2H involving the constructs detected in the first screen [38,49]. In our case, we implemented an alternative validation approach in which we investigated the correlation between interactions involving complete proteins and their domains, essentially challenging the screen. Interestingly, only interactions whose initial prey fragment highly covered the complete protein (>74%) were positive in the new assay. However, it should be noted that the unconfirmed interactions by this method should not be excluded from biological relevant interactions. Indeed, although some interactions that require full-length proteins could be missed [55], the use of protein fragments can increase the sensitivity of the screening as shown in Yang *et al*. [47], where some known interactions were only detected when domains or domains vs. full-length proteins were assayed. Likewise, the previously described interaction between LexA and P7 [29] was detected in our screen with a fragment covering 42% of the protein. Therefore, notwithstanding the limitations of the fragment-based Y2H, our results indicate that the use of a random fragment library not only expands the interaction space, but also can be useful for domain interaction determination and additionally provides a protein length coverage cutoff criteria.

### The Bam35-Bt Y2H interactome reveals the clustering of special vertex proteins and a wide modulation of host cell metabolism

Betatectiviruses have a particular lysogenic cycle, as plasmidial prophages, which raised many questions about their evolution and maintenance [66,67]. Disclosing the virus-host interactions at the molecular level would shed some light onto this growing group of enigmatic viruses and their evolutionary synergy with relevant human and animal pathogens.

After successive filtering steps, our virus-host predicted interactome contains 182 candidate interactions selected in terms of enrichment and specificity as well as biological meaning. Thus, some common patterns and interesting prey partners can be highlighted. In previous works, structural proteins were usually not involved in host-virus interactions as they are mainly involved in most intraviral interactions [38]. Conversely, a high proportion of the detected Bam35-*B. thuringiensis* PPIs are involved in Virion structure & DNA packaging. Among them, proteins located in the special vertex and membrane proteins, which are specific to tectiviruses, seem to closely interact with the host proteome. When compared with previous works on phage-host interactomes, functional annotation frequencies of the interacting host partners do not share similar patterns (Figure 6). Strikingly, although a higher frequency in protein processing and gene regulation has been associated for phage lambda with its lysogenic state, these categories are not specifically represented for Bam35 and Giles phages, which are also temperate. Nevertheless, we did detect an enrichment in proteases (see below), known to be important for phage lambda, among the Bam35-host interactions although these were not classified in the category of protein processing. Previous phage-host research mainly identified host proteins involved in transcription, replication, recombination, and repair functions. In turn, the phage proteins in the Bam35-host and Giles-host Y2H interactomes highly target host metabolic processes and transport proteins. Interestingly, despite the high differences both in the virus and the host, the Giles-host interactome approach is the most similar to ours since it is based on a host genome fragments library and Giles is also a temperate phage [41]. In any case, comparisons among different phage-host systems and techniques must be carefully taken. In line with our results, a high-thoughput proteomic analysis of the effect of the betatectivirus-related plasmid pBClin15 on its host revealed a significant impact of pBClin15 on different pathways of central metabolism during growth [68]. Similarly, diverse RNA-seq analyses have highlighted the influence of phages in membrane proteins, transporters, and metabolism [69–72]. On the other hand, some host interactors detected in our study are similar to those found in other phage-host interactomes, including ABC-transporters, NADH-quinone oxidoreductases, primosomal proteins, and phosphate regulatory proteins [38,40,41].

**Figure 6.**
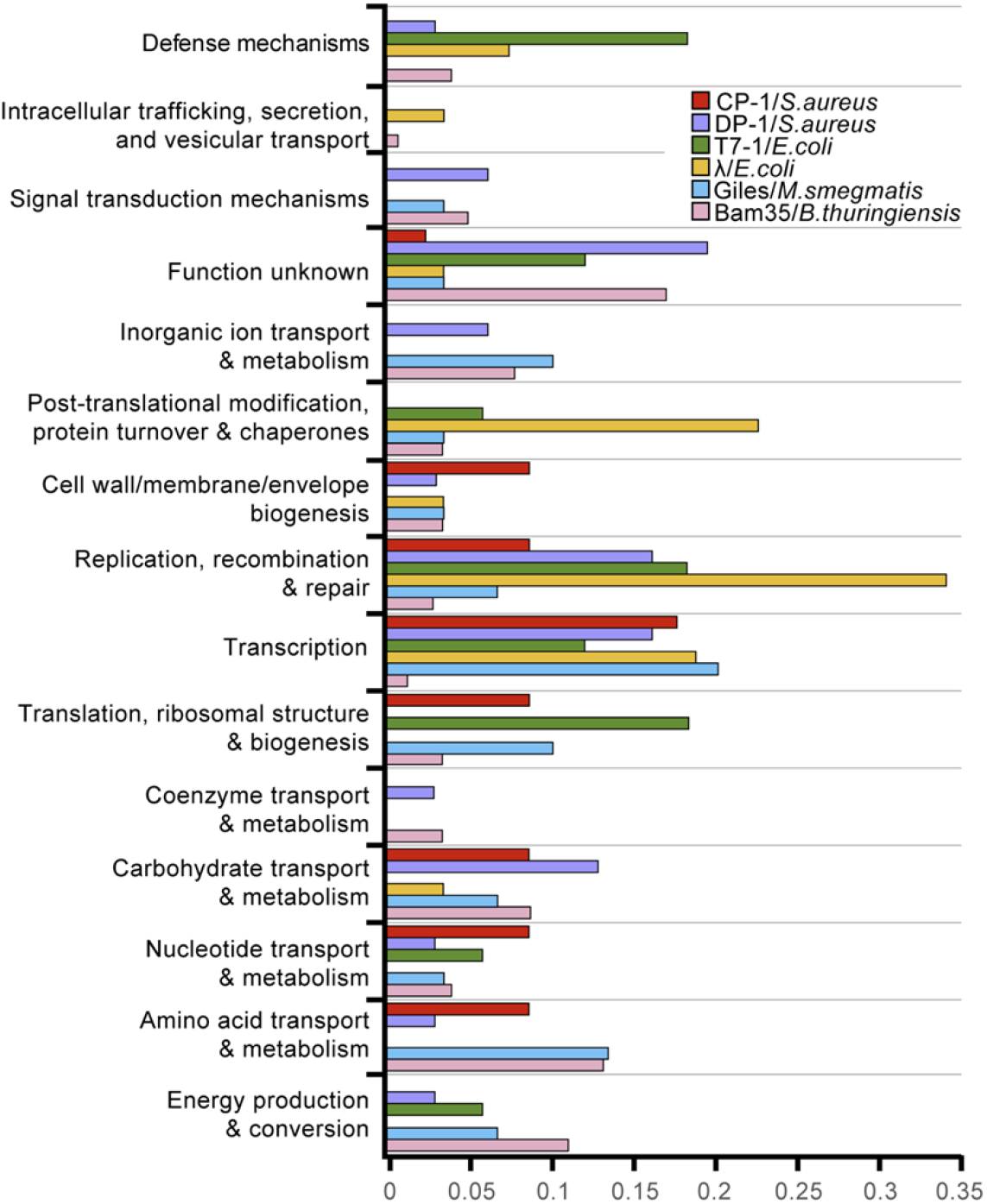
Frequency of phage-targeted proteins and their functional classes adapted from Mariano *et al*. [40]. The interactions between Giles and *Mycobacterium smegmatis* (blue) [41] and the interactions detected in this work between Bam35 and *B. thuringiensis* (pink) were added.

Among the 182 different interactions, 12.6% involved different types of peptidases. PepA (locus HBA_24805) is linked to the three functional groups of viral proteins (Supplementary Figure 5). This protein presumably functions in the processing and regular turnover of intracellular proteins, which might be subverted during the viral infection. This could also be the case for a putative M42 metalloprotease (locus HBA75_23575, also annotated as PepA) and the aminopeptidase AmpS, which interact with Bam35 structure and lysis proteins. Accordingly, *Escherichia coli* gene expression analysis during bacteriophage PRD1 infection showed that many proteases were highly induced during virion assembly [69]. Interestingly, the orthologs of these three peptidases in *Bacillus subtilis* interact with each other (String database). On the other hand, the *E. coli* ortholog of PepA (locus HBA_24805) has a role in transcription and recombination and therefore could also be related to the Bam35 life cycle regulation since PepA interacts with the viral transcription factor P6. [73]. In line with this, proteases are involved in lambda-*E. coli* interactions and linked to the viral cycle regulation, although those interactions could not be detected by Y2H probably due to their weak and transient nature [38,42]. Another protease found in the interactome, PepD, was reported to negatively affect biofilm formation [74]. Therefore, the detection of PepD as a host target, as well as a spore coat protein, could be linked to the influence of Bam35/GIL01 lysogeny on the *B. thuringiensis* sporulation rate and biofilm formation [30].

The Bam35 membrane proteins P25 and P26 interact with many host proteins which, at the same time interact with other viral proteins (Figure 5). P25 is the second most abundant protein in purified Bam35c particles [12] and P26 is thought to act as a membrane scaffolding protein [20]. As such, they could act as hubs in the interaction host-virus recruiting necessary proteins for the viral particle.

A number of viral proteins were clustered in the Bam35-host interactome by their common interactions with numerous bacterial nodes (Supplementary Figure 6). The suggested function for four of them (P15, P16, P19, and P22) is associated to the special vertex [12,20]. The clustering of these proteins in a similar host-phage interaction environment supports their proposed function. On the other hand, the localization of PRD1 P15, the Bam35 lytic enzyme P26 counterpart, in the special vertex is controversial [75,76]. Interestingly, in Bam35, our phage-host interactome suggests that P26, along with the proteins of unknown function P23 and P27 would be part of the special vertex cluster (Supplementary Figure 6). P23 has a transmembrane domain and almost all its interaction partners include different transporter proteins, hinting towards a role as membrane-associated protein related to the special vertex. P27 is also a transmembrane protein and was suggested as the penton protein [13], although the intraviral interactome results have downplayed this possibility [20]. The non-transmembrane variant of P27 only interacts with a host N-acetylmuramoyl-L-alanine amidase, a protein that degrades the cell-wall peptidoglycan. This host protein interacts at the same time with P24, proposed as the penton protein, that is also related to P30, the transglycosylase presumably responsible for the viral entry mechanism associated with the spikes [20,22]. Thus, P27 may be an anchoring virion protein whose function is related to both viral entry and release. These results could also suggest the recruitment in the virion of an additional host amidase that would help in the viral entry. Another interesting putative partner of P24 is a GroEL chaperonin, described to assist the insertion of PRD1 proteins in the virion membrane and essential for PRD1 assembly [77]. *E. coli*-PRD1 RNAseq data showed high expression levels of the host GroEL during assembly while in the case of pBClin15b, which displays cryptic prophage behavior, *GroEL* was downregulated [68,69]. Thus, it is tempting to speculate that this chaperone may also have a role in the assembly of Bam35.

Importantly, all the proteins clustered in the “special vertex group” have a predicted transmembrane domain [12] whereas the variants lacking this domain did not appear in this group. This could explain their interaction with the same partners, which are mostly membrane related transporters and may help the phage to attach to the host membrane. Particularly, the strong interactions with a putative phage tail tape measure protein from the putative prophage pLUSID3 (30985 in Supplementary Figure 6) could be due to the protein region covered by the Y2H fragment which comprises a predicted transmembrane domain. This domain appears specific to *Bacillus* phages and therefore would suggest interactions between elements of the mobilome. However, this protein is puzzling as *Caudovirales* lack any membrane within their viral particle. It should also be noted that the highly-connected P25, which also contains a transmembrane domain (as well as P32, P20, and P21), are not connected to this cluster, decreasing the possibility of non-specific hydrophobic interactions. Besides, six out of the seven confirmed interactions belong to this cluster (Supplementary Figure 6). Remarkably, within those, the enterotoxin/SH3 domain protein (Iap in Supplementary Figure 6) was also annotated as the conserved virulent factor EntA [56,78]. SH3 domains are known to be involved in protein-protein interactions and cell wall recognition and binding [79]. Since the transglycosylase P26 does not possess any signal peptide [22], it is also possible that the enterotoxin can be recruited by the phage to reach the membrane and the cell wall during and contributing to lysis.

One of the most relevant interactions detected with our approach is the described direct interaction between the viral P7 and the host LexA, key for maintaining lysogeny [19,28]. Here, the LexA fragment comprises the second half of the protein (residues 121 to 206) suggesting that the interaction domain is located in this region and it is sufficient to establish the interaction. Importantly, the detection of this previously known (and unique) interaction between Bam35 and *B. thuringiensis* provides confidence in our method. Only another viral protein interacts with the host LexA, P8. Interestingly, the ORF8 is located in a highly variable region of tectiviruses whose ORFans may alter phage regulatory functions, influencing phage and possibly also host fitness [80]. However, the lack of sequence similarity does not provide any hint about its specific function. Since the only detected host partner of P8 is LexA, it is tempting to suggest a role of P8 in viral cycle control. Particularly, as the ORF8 is located at the end of a gene cassette responsible for maintaining the lysogenic cycle [19,28], the P8-LexA interaction might also contribute to the fitness and regulation of lysis-lysogeny switch.

Although several interesting interactions have been detected, undoubtedly some may be missed by the establishment of a threshold, as shown in Yang *et al*. [47] and false positives can arise for the same reason. The knowledge of literature-based interactions has proven to be key in this matter [49]. Since our approach is the first one in the tectiviruses-host interactions field, we have combined the use of several quality filters with the previous biological knowledge of Bam35-*Bacillus* model and previous virus-host studies. Therefore, the high-throughput Y2H screening of Bam35-*B. thuringiensis*, including a high number of host proteins and tested interactions, has resulted in the detection of multiple potentially interesting and novel PPIs. Although Bam35-*B. thuringiensis* interactions need to be further analyzed with other PPIs analysis techniques and their biological relevance remains to be explored further, they open the door to new types of host-phage interactions and a deeper understanding of Bam35 and tectiviruses.

## Materials and Methods

### Nucleotides and DNAs

DNA oligonucleotides were purchased from IDT (Coralville IA, USA).

Entry donor vectors pDONR/Zeo and pCR™8/GW/TOPO™ were purchased from Invitrogen. Yeast two-hybrid system (Y2H) expression vectors pGADCg, pGADT7g, pGBKCg, and pGBGT7g [59] were available in-house.

### Bacterial and yeast strains

*Bacillus thuringiensis* HER1410 strain was originally obtained from the culture collection of the Félix d’Herelle Reference Center for Bacterial Viruses of the Université of Laval (https://www.phage.ulaval.ca) and can be retrieved with Host HER number 1410.

*E. coli* TOP10 was used for genomic HER1410 libraries generation. *Saccharomyces cerevisiae* prey strain Y187 (*MATα, ura3-52, his3-200, ade2-101, trp1-901, leu2-3, 112, gal4Δ, met, gal80Δ, URA3::GAL1*_*UAS-*_ *GAL1*_*TATA-*_ *lacZ*) and bait strain AH109 (*MATa, trp1-901, leu2-3, 112, ura3-52, his3-200, gal4Δ, gal80Δ, LYS::GAL1* _*UAS*_*-, GAL1* _*TATA*_*-, HIS3, GAL2* _*UAS*_*-, GAL2* _*TATA*_*-, ADE2, URA3::MEL1* _*UAS*_*-, MEL1* _*TATA*_*-lacZ*) were used for the Y2H screen [81].

### Genomic library construction

To generate a genomic library of *B. thuringiensis* HER1410 a four-step procedure was followed as illustrated in Supplementary Figure 1.

#### I. Generation of the genomic library and cloning into the donor vector

The genomic DNA (gDNA) of *B. thuringiensis* HER1410 was isolated using the DNeasy Blood and Tissue kit (Qiagen) and concentrated by ethanol precipitation [82]. The extracted gDNA was partially digested with the restriction enzyme CviAII (New England Biolabs) and subsequently 5’-dephosphorylated with Calf intestinal alkaline phosphatase (New England Biolabs). CviAII cuts on a sequence motif ‘CATG’ which is highly frequent on bacterial genomes, with 17,370 sites in HER1410 genome, cutting on average every 356.8 base pairs (bp). The digested DNA was separated by agarose electrophoresis and an agarose block with DNA fragments spanning between about 450-750 base pairs (marker bands) was cut-out, subjected to gel-extraction DNA purification with QIAquick Gel Extraction Kit (Qiagen) and purified by ethanol precipitation [82]. Gap filling of the 3’ overhangs and A-tailing were performed using Taq DNA polymerase. The library was subsequently cloned into the plasmid pCR™8/GW/TOPO™using the pCR™8/GW/TOPO™ TA Cloning Kit (Thermofisher).

#### II. Amplification of the genomic library in pCRTM8/GW/TOPO®

The donor plasmid library was transformed in *E. coli* Top10 cells (CBMSO Fermentation Facility) by electroporation. Transformants were stored at −80 °C in the presence of 10% (v/v) glycerol. The number of independent clones was determined as CFU on LB-Agar plates supplemented with 100 µg/ml spectinomycin. According to the expression given by Clarke and Carbon [83], the colony bank size needed to obtain a plasmid collection that covers *B. thuringiensis* HER1410 genome at least once, with a probability of 0.95 and an average fragment size of 600 bp, taking into account the six possible reading frames, is 1.842 × 10^5^ CFU. This number of clones was outreached along the construction of the library (Supplementary Figure 1). The original genomic library was propagated in SeaPrep® (FMC) semisolid medium [84] to minimize representational biases that can occur during the expansion of plasmid DNA libraries [85]. The total number of clones was titered again by serial dilutions and plating on selective LB-Agar plates. Then, 10^7^ cells were inoculated in 50 ml LB medium supplemented with 100 µg/ml spectinomycin and grown overnight at 37 °C for purification of the plasmid library DNA by miniprep (NucleoSpin Plasmid, Macherey-Nagel). The efficiency of fragment insertion (100%) was checked by plasmid purification of 15 clones followed by digestion with EcoRI-HF (Supplementary Figure 7A) and sequencing (Supplementary Figure 7C) with the forward primer GW1 (Supplementary Table 7).

#### III. Subcloning of the library in the Y2H expression vectors

Using 50 ng of the pCR8/GW/TOPO library as donor plasmids, the inserts were subcloned into the pPC (pGADCg) and pPN (pGADT7g) plasmids using Gateway™ LR Clonase™ II Enzyme mix (Thermofisher), according to the manufacturers’ instructions. The LR reaction product was used to transform Top10 cells, titered, and amplified as detailed above (Supplementary Figure 1). LR reactions with pPC and pPN result in C-terminal and N-terminal fusions of the Gal4p activation domain (AD) to the gateway cassettes respectively. The efficiency of insertion (85% for pPC and 100% for pPN) was checked by plasmid purification of seven clones from each library followed by PCR amplification (Supplementary Figure 7B). These fourteen clones were also verified by sequencing (Supplementary Figure 7C) using attB1 and T7_FW primers respectively (Supplementary Table 7). The genomic libraries in pPC and pPN prey vectors were purified by miniprep as detailed above.

#### IV. Transformation of *Saccharomyces cerevisiae* Y187 with the Y2H genomic libraries

pPC and pPN libraries were used to transform *Saccharomyces cerevisiae* strain Y187 by electroporation [86]. Transformants were titered to determine the number of independent clones (Supplementary Figure 1) and the library was further expanded in 50 150-mm Petri dishes with selective solid media without leucine. Grown cells were harvested in liquid selective medium containing 25% glycerol, for storage at −80 °C. Prior to mating experiments, the final libraries were titered on selective media, and their genome coverage was analyzed by high-throughput sequencing (see below).

### Yeast two-hybrid screening

We designed a multiple yeast two-hybrid screening with a collection of 80 bait vectors previously characterized in Berjón-Otero *et al*. [20] that include C-terminal (pBC (pGBKCg)) and N-terminal (pBN (pGBGT7g)) fusions of the Gal4p DNA-binding domain (DBD) to all the 32 Bam35 ORFs plus six variants without terminal transmembrane domains, as well as the empty pBC and pBN vectors. Each of the bait vectors was tested against the prey HER1410 libraries in pPC and pPN vectors, yielding a total number of 158 mating experiments. Since it was not possible to obtain the Bam35 ORF16 pPC construct, this vector was not included in the screen [20]. Mating experiments were performed between AH109 yeast cells harboring pBC or pBN with the Bam35 ORFs (bait) and Y187 yeast cells harboring the pPC or pPN libraries as described in Mehla *et al*. [87], with specific modifications for the subsequent HTS analysis (see next section). Briefly, 4 ml of bait and prey cultures at an OD_600_ of 0.8-1 were mixed before harvesting the cells by centrifugation. Cells were subsequently plated on rich solid media (YPDA plates) to allow mating. After overnight incubation at RT, cells were harvested, washed, and resuspended in 2 ml of selective liquid media lacking leucine, tryptophan, and histidine (-LWH). To perform the interaction selection, *HIS3*, coding for an imidazole glycerol phosphate dehydratase necessary for histidine biosynthesis, was used as a reporter gene. In short, 100 µl of the resuspended cells were plated on selective solid media (-LWH) supplemented with 3-amino-triazole (3AT). To avoid self-activation by DBD-fusion proteins, each screen was performed in the presence of the 3AT concentration determined for each bait [20]. To measure the efficiency of mating, the culture was diluted 1:10^5^ and plated on diploid selective solid media without leucine and tryptophan. Moreover, to remove the background and further confirm the positive clones, we made two stamp replicas on triple dropout media in the presence of X-α-Gal. Blue colonies show positive interactions for two markers, *HIS3* and *MEL1*. In some cases, the screen gave rise to a lawn of cells. 3AT was increased for those screens to reduce the number of false positives (Figure 3A). Finally, Y2H-positive colonies i.e. yeast cells capable of growing on the triple dropout media, from the last replica of each Y2H screen were pooled and harvested for further HTS analysis. Additionally, to validate HTS results (see below), we randomly picked single colonies of the final replicas before harvesting. As such, we randomly analyzed 70 colonies including all combinations (pBC-pPC, pBC-pPN, pBN-pPC, pBN-pPN) whose prey inserts were identified by colony PCR followed by Sanger-sequencing (Supplementary Table 9).

### Genomic library and Y2H positives analyses by high-throughput sequencing

#### pPC and pPN libraries sequencing

Validation of the HER1410 genomic libraries was achieved by sequencing each of the plasmid libraries, pPC and pPN, purified from the yeast strain Y187 with the Zymoprep Yeast Plasmid Miniprep kit (Zymo Research). Plasmid isolation from yeast cells typically yields very low amounts of DNA. As such, the purified plasmid libraries were used to amplify the genomic inserts by tailed-PCR (Supplementary Figure 7D) with the forward primers attB1_HTS for pPC library and T7_HTS for pPN library and the reverse primer attb2_HTS (Supplementary Table 7). To ensure faithful and proportional amplification of each fragment, Q5 High-Fidelity DNA Polymerase (New England Biolabs) was used, and the number of amplification cycles was limited to 20. Secondary PCR amplification was subsequently performed by the HTS Unit from the “Parque Científico de Madrid” to generate Illumina sequencing libraries. PCR products were sequenced using a single Illumina MiSeq 300-bp paired-end run, in this facility. The total raw reads obtained from the libraries sequencing would represent an average coverage of the original genome (6,147,475 bp) of 260X for pPC-library and 234.5X for pPN-library. Prior to the construction of Illumina sequencing libraries, fragments were analyzed by agarose electrophoresis. Illumina sequencing libraries were analyzed by microchip electrophoresis on an Agilent 2100 Bioanalyzer using DNA 7500 Assay Kit (Agilent Technologies).

#### Y2H positives preparation and sequencing

The Y2H-positive colonies from the last replica of each Y2H assay were pooled and harvested. This resulted in 156 samples, one per mating experiment. To identify the interacting prey partners for each sample, plasmid DNA was purified by yeast miniprep in multi-well MW96 format (Zymoprep-96 Yeast Plasmid Miniprep). Then, the HER1410 fragments were low-cycle, tailed PCR-amplified using Q5 High-Fidelity DNA Polymerase, as explained above, with the specific forward primers pPC and pPN and the reverse primer attB2_HTS (Supplementary Table 7). Moreover, each sample was confirmed by PCR with specific oligonucleotides for the corresponding viral bait gene. Finally, PCR products were sent to the HTS facility to construct barcoded Illumina libraries, that were sequenced in a single Illumina MiSeq 300-bp paired-end run to obtain about 15,000 reads per sample. Fragments were analyzed by agarose electrophoresis prior to the construction of Illumina sequencing libraries. Illumina sequencing libraries were analyzed by microchip electrophoresis on an Agilent 2100 Bioanalyzer using DNA 7500 Assay Kit (Agilent Technologies). The samples containing fragments shorter than 300 bp were split (<300 bp and >300bp) and sequenced independently to avoid sequencing bias.

#### Trimming, quality check, and mapping of Illumina reads

The Illumina reads were verified for quality using FastQC v0.11.8 and Trimmomatic v0.38 was used to exclude reads shorter than 185 bp [88,89]. To remove primers and plasmids sequences from the reads, a clipping step based on sequence size was performed using seqtk [90]. The first 39 and 139 nucleotides of R1 reads of pPC and pPN libraries respectively, and the first 38 nucleotides of R2 reads were removed. For seqtk clipping of the Y2H positive sequences, the first 99 and 59 nucleotides of R1 reads from pPC and pPN sequences respectively, and the last 38 nucleotides of R2 reads were removed. FastQC reports of raw and processed reads were consolidated using MultiQC v1.9 [91]. To extract the in-frame HER1410 fragments, a custom bioinformatics pipeline was set up (Figure 1). Briefly, reads that did not start with the starting codon “ATG” were removed. These “ATG” sites correspond to the prey fragment ends generated by partial digestion with CviAII, cloned in frame with the destination vectors. Subsequently, using R1 and R2 reads, prey fragments were reconstructed and clustered generating a database of unique fragments. Clustering was performed using CD-hit software at 100% sequence identity level and identical length [92]. The unique fragments were then translated and searched in the HER1410 proteome using BLASTp [93]. Fragments resulting in a BLASTp hit longer than ten amino acids were kept as “in-frame unique fragments”. To obtain the frequency for each unique fragment, clusters reconstruction was performed resulting in the number of reads for each unique fragment. Additionally, results were manually curated to eliminate the fragments containing a stop codon or that were not in frame with the AD domain in pPC, and the hits after a stop codon in pPN, generating our Y2H-validated fragments database. The bioinformatics pipeline is available on github: https://github.com/LoGT-KULeuven/y2h_Bam35-Bt_analysis

#### Evaluation of genomic library quality

Genomic library statistics were analyzed using Excel software (Microsoft). The nucleotide coverage of total fragments was calculated using bwa-mem [94], samtools [95] and visualized with weeSAM version 1.5 (accessed at https://bioinformatics.cvr.ac.uk/weesam-version-1-5/).

#### Evaluation of raw Y2H interactions

The mapping of positive interactions reads resulted in the identification of 4,477 possible interactions (Supplementary Table 3). Filtering of these raw results was performed to improve the specificity of the putative protein interaction set by applying a set of sequential filtering steps to the data. First, the number of reads corresponding to each interaction was normalized to the total reads in the sample. This normalized number was used as a measure of enrichment of the prey fragment in the corresponding combination and, therefore, of the strength of the interaction. Thus, this value was used to establish enrichment categories according to the distribution of the positive hits (Supplementary Figure 10). The dataset was divided into three categories: C for interactions with 0-0.25% of abundance, B for 0.25-10%, and A for 10-100%. Only the most abundant interactions (categories A and B) were considered for further analysis. Second, prey fragments interacting with a large number of baits were tagged as potential promiscuous or “sticky preys” (Supplementary Table 4). Thus, interactions involving HER1410 protein fragments that interact with more than six different Bam35 proteins, the average of interactors after selection, were excluded from further evaluation. Third, prey interacting partners of bait pBC and pBN empty plasmids were considered as false-positive generating preys. Interactions involving these prey plasmids were also removed from the dataset (Supplementary Table 5). Last, duplicated interactions coming from under and over 300 bp sequencing runs were consolidated. Finally, the obtained putative PPIs resulted from dataset filtering were analyzed and represented as interaction maps with Cytoscape software [96].

Descriptive and comparative analysis of the positive interactions reads dataset (Supplementary Table 2) was performed with SPSS® Statistics software (IBM). Linear correlation between library fragments and interacting fragments reads abundance was performed with the ggplot package for R software [97].

### Full-length protein pairwise Y2H assays

A total of 33 putative interactions were re-screened using complete host proteins and their viral partners (Table 3). In total, 15 interactions from the category A and 18 interactions from the category B were selected at random and retested with the full-length host proteins. The *B. thuringiensis* HER1410 purified genomic DNA was used to amplify the selected ORFs by PCR using Q5 High-Fidelity DNA Polymerase and ORF specific primers (Supplementary Table 7). These primers were designed using Geneious software [98], ensuring the presence of 20 to 30 nucleotides complementary to the ORF of interest and removing endogenous stop codons. The *attB1* and *attB2* sequences were added at the 5’ end of forward and reverse primers, respectively. As described in Berjón-Otero *et al*. [20], tail PCR products were cloned into the entry vector pDONR/Zeo (Invitrogen) and subsequently subcloned into the corresponding prey vectors pPC and pPN. All vectors obtained from cloning and subcloning steps were tested by colony PCR with ORF specific primers. The insertion of the ORFs into the entry and expression vectors was also verified by sequencing. The expression vectors were used to transform *S. cerevisiae* Y187 following the heat-shock transformation protocol described in Mehla *et al*. [87]. Transformants containing prey vectors were selected on solid medium without leucine and checked by colony PCR using pPC_HTS or pPN_HTS forward primers and the corresponding ORF specific reverse primer (Suppementary Table 9).

Pairwise screens of the selected interactions were performed, as detailed in Table 3, using the bait vectors pBC and pBN containing the Bam35 ORFs characterized in Berjón-Otero *et al*. [20] and the obtained pPC and pPN prey vectors containing the selected HER1410 ORFs. We performed mating between yeast cells containing bait vectors and cells containing prey vectors on rich solid media (YPDA plates) including the correspondent self-activation controls with the empty vectors. Diploid cells were selected on solid medium without leucine and tryptophan. Finally, to detect protein-protein interactions, *HIS3* was used as reporter gene. For this, the obtained diploid cells were plated on selective solid media without leucine, tryptophan, and histidine supplemented with different 3AT concentrations (0, 0.025, 0.1, 3, 10, 25, 50, and 100 mM). When the yeast growth was higher in the Bt-B35 combination than in the self-activation controls, this interaction was considered as positive.

## Data availability statement

All sequencing data was deposited in the NCBI SRA database and is accessible via the BioProject PRJNA717632, or directly via the sample accession numbers listed in Supplementary Table 2. The Bam35-*B. thuringiensis* high-quality interactions reported in Supplementary Table 6 have been also deposited in the Database of Interacting Proteins (https://dip.doe-mbi.ucla.edu/) under the IMEx Consortium dataset identifier IM-28933.

## Acknowledgments

This work was funded by grant from Fundación Ramón Areces [VirHostOmics]. A. L. was holder of a PhD fellowship [FPU15/05797] from the Spanish Ministry of Science, Innovation and Universities. C.L. is supported by a PhD fellowship from FWO Vlaanderen (1S64720N).

We would like to thank the Genomics and Massive Sequencing facility of CBMSO for technical assistance at the early stages of this project. An institutional grant from Fundación Ramón Areces and Banco Santander to the Centro de Biología Molecular Severo Ochoa is also acknowledged.

